# Incorporating Randomness into DNA Steganography to Realize Secondary Secret key, Self-destruction, and Quantum Key Distribution-like Function

**DOI:** 10.1101/725499

**Authors:** Meiying Cui, Yixin Zhang

## Abstract

DNA has become a promising candidate as future data storage medium, which makes DNA steganography indispensable in DNA data security. While PCR primers are conventional secret keys in DNA steganography, the information can be read once the primers are intercepted. New steganography approach is needed to make the DNA-encoded information safer, if not unhackable. Herein, by mixing information-carrying DNA with partially degenerated DNA library containing single or multiple restriction sites, we build an additional protective layer, which can be removed by desired restriction enzymes as secondary secret keys. As PCR is inevitable for reading DNA-encrypted information, heating will cause reshuffling and generate endonuclease-resistant mismatched duplexes, especially for DNA with high sequence diversity. Consequently, with the incorporation of randomness, the DNA steganography possesses both quantum key distribution (QKD)-like function for detecting PCR by an interceptor and self-destructive property. With a DNA-ink incorporating the steganography, the authenticity of a writing can be confirmed only by authorized person with the knowledge of all embedded keys.

---

As a novel data storage medium, DNA possesses high capacity and longevity, and can be amplified and operated biochemically. The remarkable technological improvement in *de novo* DNA synthesis has made the use of synthetic DNA as data storage medium feasible^1,2^. Recently, encoding digital data into DNA sequences and retrieving the original file without errors have been reported by several research groups^3–9^. However, like every other data storage media, communication with DNA can be intercepted and copied, which has made DNA steganography an important field in DNA data security. DNA steganography provides protective layers to DNA-encrypted data by mixing dummy DNA sequences with intended information DNA. Clelland et al. have for the first time turned DNA steganography into reality by hiding information-carrying DNA in human genomic DNA fragments and spotting them as microdots on filter paper^10^.

Currently, most DNA steganography methods utilize primer sequences as secret keys, which are shared secretly between the sender and recipient to maintain a private information link^11^. The more complex the keys (e.g., more and longer keys), the securer the encrypted information ^12, 13^. However, such secret keys are breakable upon subjecting the intercepted information to intensive analysis. In recent years, quantum key distribution (QKD) has been suggested to be able to provide an additional layer of protection for the existing encryption algorithm, as its unique feature is the ability of the two communicating users to detect the presence of any third party^14,15^. However, currently the rate-distance limit of QKD hinders large-scale deployment of QKD network ^16-18^.

Can we realize QKD-like function in DNA steganography utilizing some superior intrinsic properties of DNA over qubits? While QKD is based on the feature that the process of measuring a quantum system disturbs the system, is there a technical analogue of such “measurement” in DNA analysis? Although there are many methods to sequence DNA, for a sample of information-carrying DNA in small quantity, PCR will be inevitable for the interceptor (“Eve”) to test different keys or key combinations, to amplify the sample for sequencing, as well as to make copy for the intended recipient (“Bob”). Therefore, if we can make the PCR process generate a disturbance and leave a trace, the communicating users will be able to detect the presence of the third party, analogues to the QKD-based communication. If the method can be designed based on a physical principle as fundamental as quantum superposition (for QKD), the resulting DNA steganography will be extremely safe, if not unbreakable.

Here we report the experimental implementation of DNA steganography method, integrating QKD-like function, a secondary secret key, and a self-destruction mechanism. Through mixing information-carrying DNA (i-DNA) with a partially degenerated DNA library (d-DNA), the randomness in sequences provides not only an additional mask (e.g. as using genomic DNA) to cover the encrypted information, but also a heat-induced reshuffling mechanism to generate mismatches during the re-annealing steps of PCR. It is important to note that the increase of entropy associated with hetero-duplex formation through reshuffling is thermodynamically favorable. Thus, without the right combination of restriction enzyme pre-treatment and primers (**Fig. 1**), heating associated with PCR amplification will leave a permanent trace (QKD-like function) and make the information unreadable, even when the sample is afterwards subjected to the right processing (self-destruction).

**Figure 1.**
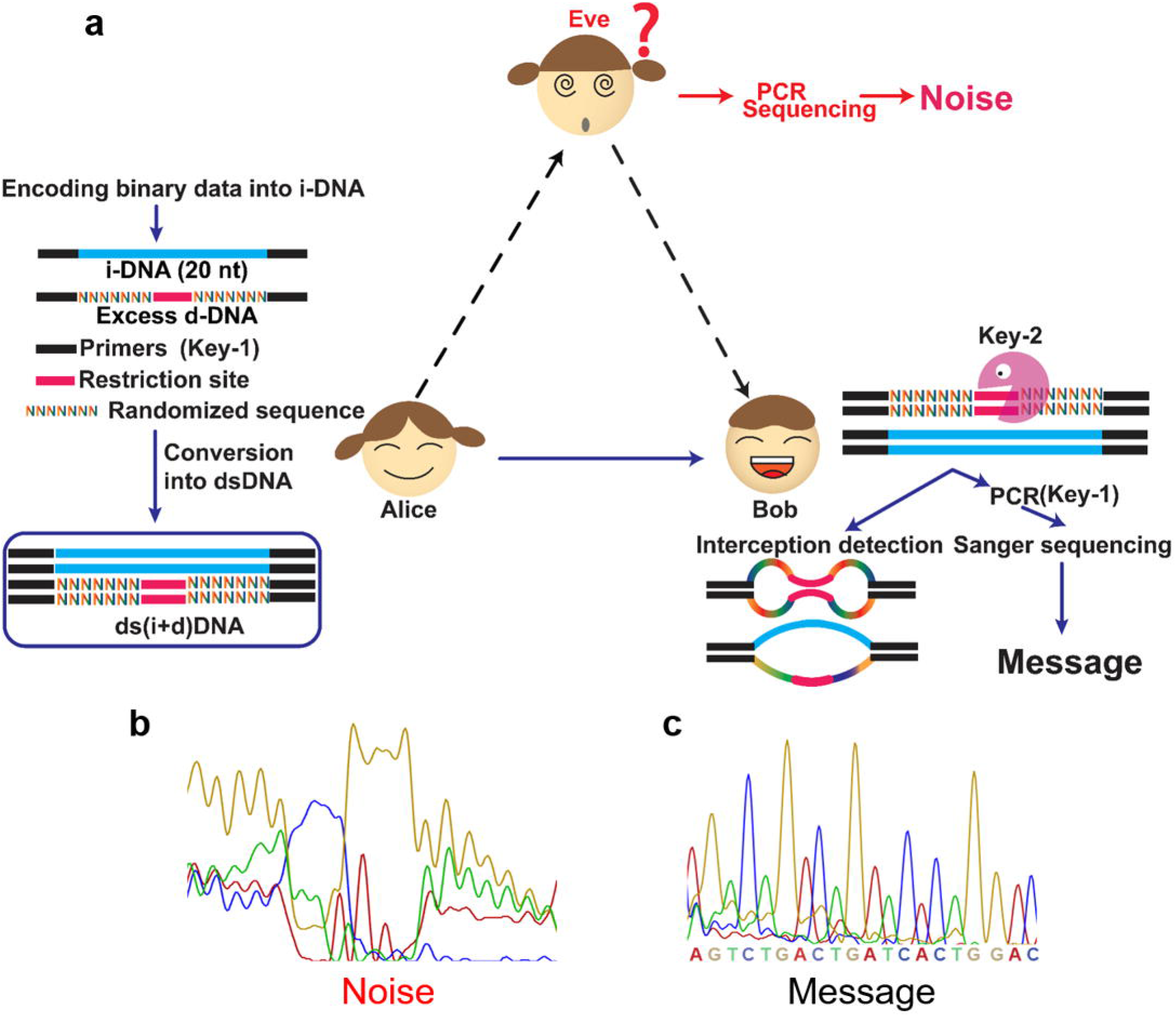
Design of DNA steganography. (a) Scheme of the proposed DNA steganography method with incorporated randomness. Alice generates an encrypted message and delivers it to Bob. Bob will use the combination of key-1 and key-2, to be able to sequence the DNA and retrieve the information, while he can also use qPCR to detect potential interception (by Eve). (b) Sanger sequencing chromatogram of the message by Eve using the correct primers (Key-1). (c) Sanger sequencing chromatogram of the message by Bob using the correct primers (Key-1) after treating the sample with the correct restriction enzyme (Key-2).

## Results

### DNA steganography design

As depicted in **Fig. 1**, sender Alice converts a binary message into a 20 nt DNA sequence using a classical substitution cipher ^19^. The conversion algorithm required to translate the binary message into a DNA sequence is not in the scope of this work. The message sequence is flanked by two primer sequences and the resulting i-DNA is mixed with a d-DNA (in large excess). The d-DNA possesses the same length and shares the same primers as i-DNA, thus can remain as a mask to cover i-DNA even when the primer information is known to an interceptor (**Fig. 1b**). The middle 20 nt-region of the d-DNA contains a 6 nt-restriction site flanked by two 7 nt-randomized sequences, representing a 4^14^ sequence diversity. The single strands were then converted to dsDNA by DNA polymerase. Alice and Bob, but not Eve, share the information regarding primers (key-1) and restriction sites (key-2). Receiving the message from Alice, Bob first digests the sample with the desired restriction enzymes (key-2), subjects the product to PCR with desired primers (key-1), and performs sequencing to read the encrypted message (**Fig. 1c**).

After intercepting the message, Eve cannot read the information without both keys. If she knows the primers (key-1), the most commonly used secret key, she will amplify the sample by PCR to obtain an adequate amount for sequencing. However, as the information in i-DNA is masked by d-DNA, Eve will not be able to distinguish the message from the d-DNA noise. As shown in **Fig. 1b**, without the restriction enzyme pre-treatment (key-2), the information cannot be read through sequencing.

In addition to key-1 and key-2, the steganography also provides a self-destructive feature and a QKD-like function. In the attempt of decoding the message, Eve is required to amplify the intercepted DNA sample by PCR. However, when amplifying a highly diverse DNA pool, mismatched dsDNA will be generated during the iterated melting and assembling cycles. The mismatches around the restriction sites will affect the substrate recognition by DNA endonucleases, diminishing the efficiency of digestion. Therefore, performing PCR prior to enzyme digestion is a self-destruction process: the mask layer can no longer be removed, as the formation of mismatches cannot be reversed due to the highly diverse d-DNA in large excess. In order to maintain the flow of the communication, after intercepting the message, Eve needs to send the sample to Bob, otherwise Bob would consider it lost and inform Alice. Eventually, Bob can use the secret keys not only to decrypt the message, but also to detect whether the message has been “read” before. After performing the restriction enzyme digestion (key-2), Bob can use quantitative PCR (qPCR) to determine the efficiency of enzyme digestion from the ΔC_t_ value and meanwhile evaluate the sequence distribution and diversity from the shape of amplification curve and melting curve ^20-22^ (**Fig. 2**). Peak at a low temperature in the melting curve indicates the presence of less stable heteroduplex, while peak at abnormally high temperature implies higher-molecular weight (MW) structures, which are produced by and accumulated over PCR cycles, especially for templates with high sequence diversity^20, 23^. With the increase of sequence diversity, the qPCR amplification curve will transform gradually from a sigmoid curve to a bell-shaped curve ^20,21^. Therefore, by qPCR measurement Bob can judge whether the DNA has been intercepted and subjected to analysis.

**Figure. 2.**
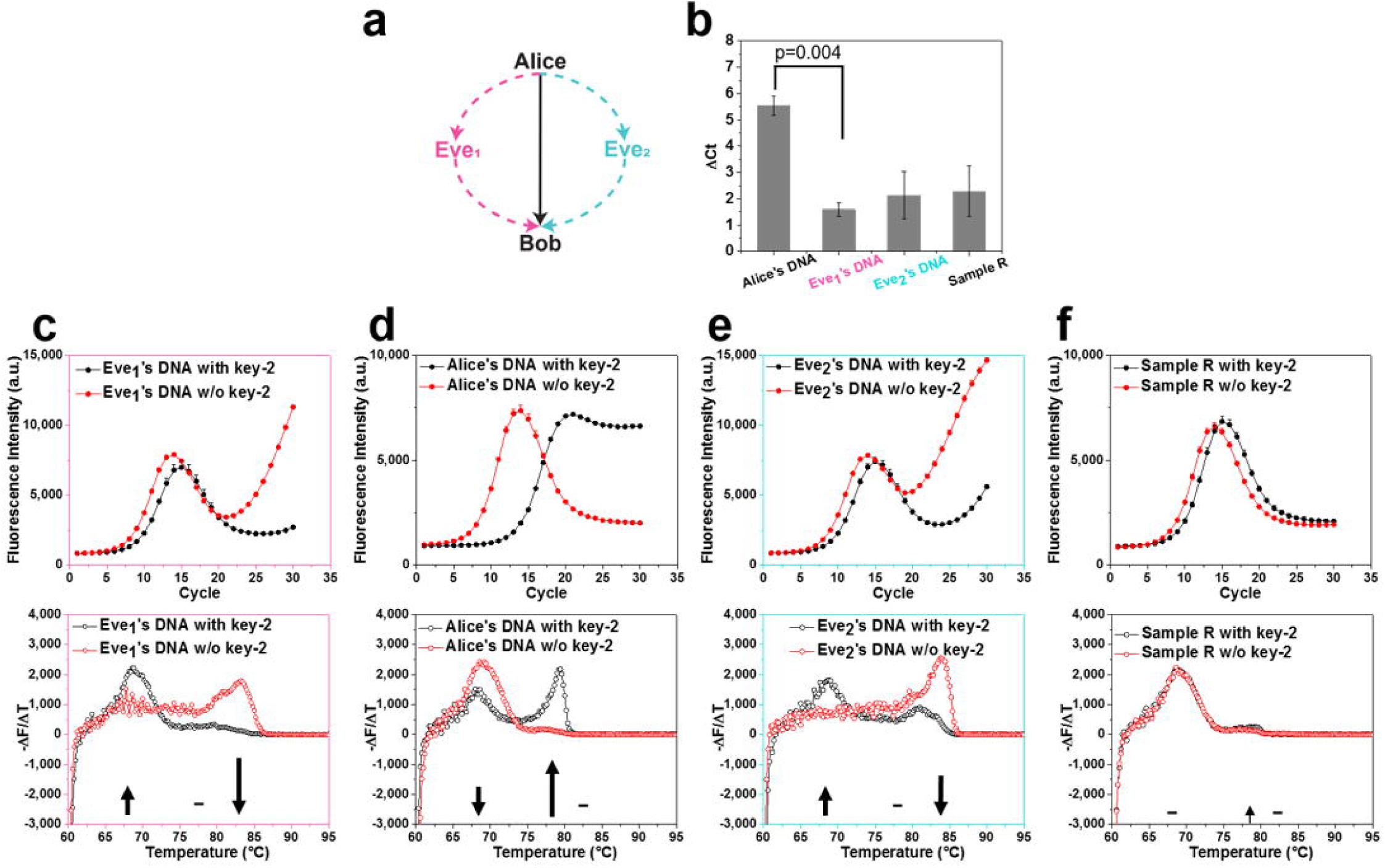
Interception detection. (a) Bob can receive three different types of sample from Alice, Eve1 (who knows the key-1), or Eve2 (who does not know the key-1), respectively. (b) ΔCt (Ct of treated sample - Ct of untreated sample) values of Alice, Eve1, and Eve2’s DNA, as well as sample R before and after key-2 treatment. Statistical significance was assessed with paired one-tail t-test. DF=2, t-value=2.92. All experiments were performed independently three times. (C-F) qPCR amplification curves (top) and melting curves (bottom) of Eve1’s DNA (c), Alice’s DNA (d), and Eve2’s DNA (e), and R (f) with and without key-2 treatment. The change of melting peak distribution is indicated by arrows, - represents no significant change before and after key-2 treatment. Error bars represent standard deviation (n=3) DF: degree of freedom

### Experimental validation

By mixing i-DNA with d-DNA, the incorporation of randomness aims to give DNA cryptography a second layer of secret key, a self-destructive feature, and a QKD-like function. We have chosen the restriction enzyme SmaI recognition site CCCGGG as the cleavage site in d-DNA, while i-DNA and d-DNA share the same primer sequences (**Fig. 1**). First, a suitable ratio of i-DNA and d-DNA was investigated. i-DNA was mixed with the d-DNA at three different ratios (1:1, 1:10, and 1:100). The samples were amplified for 25 PCR cycles, and subjected to sequencing, simulating the operations that Eve would perform upon intercepting the message and knowing the information of key-1. As shown in **Fig. 1b** and **S1**, the sequence of i-DNA can be read when i-DNA and d-DNA were mixed at 1:1 ratio, became obscure at 1:10, and was completely masked at 1:100 (Fig.1b). Therefore, 1:100 ratio was used for the following experiments.

Next, we investigated the effect of SmaI treatment on four different types of samples (**Fig. 2**): the 1:100 mixtures of i-DNA and d-DNA (i+d-DNA) (i) without a PCR pre-amplification step; (ii) with PCR pre-amplification using correct or (iii) wrong primers; and (iv) with identical thermocycling in the absence of polymerase and primers (Sample R). i+d-DNA without PCR pre-amplification represents the authentic message from Alice to Bob, while the pre-amplified sample can mimic the message from Eve, which was intercepted and amplified (using either correct key-1 (Eve_1_) or wrong key-1 (Eve_2_)). The sample R allows us to investigate the effect of heat-induced reshuffling in the absence of polymerase on restriction enzyme recognition. Four samples of the same concentration were digested with SmaI and subjected to qPCR with desired primers (key-1). In the qPCR measurement, 30 cycles of amplification and one cycle of melting procedure were performed. As shown in **Fig. 2b**, the efficiency of enzyme digestion on R, Eve_1_ and Eve_2_ were remarkably lower than that of Alice’s DNA, as evidenced by the ΔC_t_ values. Sequence distribution and diversity of the samples can be analyzed by the shape of qPCR amplification curve (**Fig. 2**). Upon SmaI treatment, i-DNA, without a cleavable site for the enzyme, became dominant in authentic sample but not in PCR pre-amplified sample, as shown by the loss of characteristic bell-shaped curve (**Fig. 2d**). Moreover, the fluorescence signal started to increase in the later cycles of qPCR in Eve_1_ and Eve_2_. Surprisingly, for Eve_2_, although the template was not amplified due to the use of wrong primers, the presence of polymerase can cause high end-point fluorescence signal in the qPCR curve, as the DNA sequences with high diversity can intertwine with each other to generate structures of high melting temperature (as shown later). In the absence of polymerase, the bell-shaped curve of sample R were not affected by the SmaI treatment (**Fig. 2f**).

The effects of SmaI treatment on different samples can also be assessed by analyzing the melting curve. After enzyme digestion, the melting peak of the authentic sample has shifted from the range of 65 - 70 °C to 75 - 80 °C, indicative of hetero- and homo-duplex dsDNA, respectively (**Fig. 2d**). Before enzyme digestion, Eve_1_’s DNA pre-amplified with correct key-1 showed a peak at 80 - 85 °C, indicating high-MW DNA structures (**Fig. 2c**). Upon enzyme digestion the melting curve redistributed into a remarkable decrease of the main peak at 80 - 85 °C and a dramatic increase at 65 - 70 °C. Eve_2_’s DNA also showed a peak at 80 - 85 °C, which decreased but remained strong after enzyme digestion (**Fig. 2e**). Enzyme digestion caused insignificant increase of the 75 - 80 °C peak for both Eve_1_ and Eve_2_, as compared to the authentic sample. Therefore, after enzyme digestion, if Bob observes a low ΔC_t_ value, a bell-shaped qPCR curve, and a melting peak at high temperature (either with or without enzyme digestion), he will be alarmed regarding the interception and possible leakage of key-1.

After treating the sample with SmaI (key-2) and PCR amplification (key-1), Bob will read the message by Sanger sequencing (**Fig. 1c**). Because enzyme digestion cannot remove the mask of d-DNA in samples previously subjected to PCR amplification, Bob can only decipher the authentic message from Alice, but not the copy generated by PCR amplification. The interception leaves a trace (QKD-like function) and destroys the message (self-destruction). We then investigated the limit of enzyme digestion to distinguish i-DNA from d-DNA. When i-DNA and d-DNA are mixed in 1:3 ratio, i-DNA can be clearly read by sequencing without using SmaI, as each base in the degenerated segment at a given position produces only 3/4 of the signal intensity as compared to that from i-DNA at this position. Therefore, if a digestion product possesses a sequence distribution similar to the 1:3 mixture, the samples can be easily read by sequencing. We compared the qPCR curve of 1:3 mixture with those from various mixtures after SmaI treatment (**Fig. S2** and **S3**). At the ratio of 1:200, but not 1:500 and 1:1000, the sequence distribution of digested product was close to that of 1:3 mixture. Sequencing results also demonstrated that i-DNA masked in the 1:200 sample, but not in the 1:500 and 1:1000 samples, could be clearly retrieved (**Fig. S4**). This experiment shows the current technical limit for masking i-DNA with d-DNA, as further increasing the d-DNA concentration will make the message unreadable for Bob.

### DNA steganography with multiple secondary keys

Next, we investigated whether other restriction site/enzyme can also be used as key-2, and whether a combination of two different restriction sites/enzymes can generate a more complex key-2. We combined two different types of d-DNA containing SmaI restriction site and EcoRV restriction site in equal amounts and mixed with i-DNA (50:50:1) to increase the complexity of the protection layer. The mixture was first treated by SmaI, and followed by EcoRV, then monitored by qPCR and Sanger sequencing (**Fig. 3**). Treatment with SmaI did not change the bell-shaped amplification curve, while the low ΔC_t_ value of 0.89±0.1 reflected that only half of the mask was removed. Remarkably, an additional digestion step by EcoRV tremendously changed the qPCR curve, indicating a dramatic change in sequence distribution and concentration (ΔC_t_ = 3.52±0.17). The message could be fully retrieved by sequencing only after treatments with both enzymes (**Fig. 3c**-**e**). Only the combination of SmaI and EcoRV can generate a functional key-2, together with the desired primers (key-1), for decrypting the concealed information.

**Figure 3.**
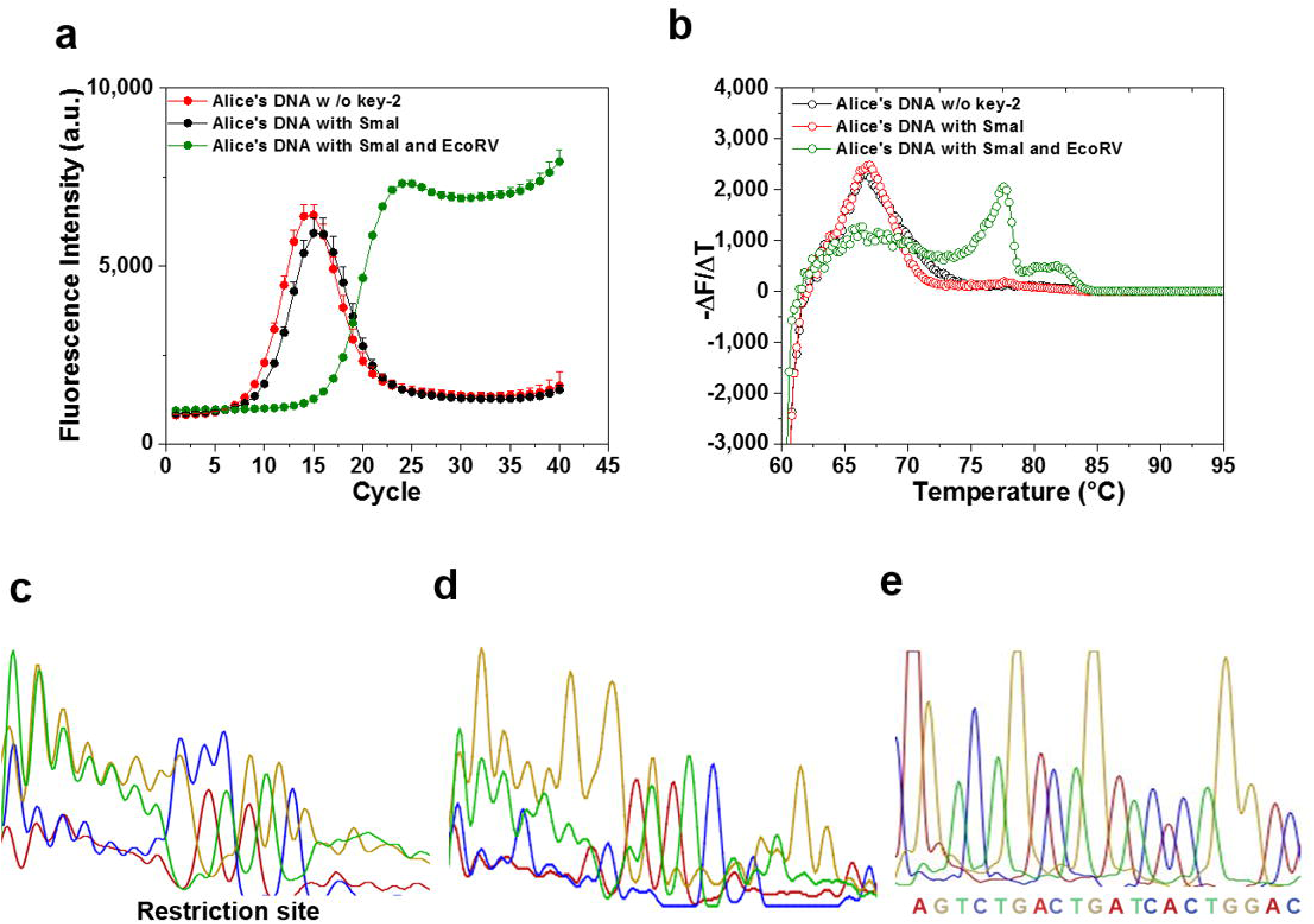
Incorporation of multiple key-2. qPCR amplification curves (a) and melting curves (b) of i-DNA concealed with two types d-DNA. Sanger sequencing chromatogram of i-DNA concealed with two types of d-DNA before key-2 treatment (c), after SmaI treatment (d), and after SmaI + EcoRV treatment (e).

Because of the fast developments in digital and engineering technologies, the traditional function of signature for self-identification has never been so severely challenged. To demonstrate that the DNA steganography can be used to produce materials for signature, whose authenticity can be confirmed only by people with the knowledge of the secret keys, the mixture of i-DNA and d-DNA was added to an ink. As shown in figure **4a**, the Chinese character “secret” (in an ancient seal script) was written on filter paper with the DNA ink. To analyze the DNA incorporated into the writing, the paper was cut into small pieces and incubated in water. The DNA in solution was then extracted with a DNA purification cartridge, and subjected to the decoding procedure using the two secret keys. As shown in figure **4b** and **4c**, only with the right keys, PCR amplification led to a sample with relatively simple composition, which can be sequenced to reveal the correct information (figure **4d**). In principle, using this methodology, it is possible to further enhance the security of steganography by using multiple secondary keys in order to embed basic logic operators into the DNA-encrypted message. For example: 1) multiple restriction sites in different d-DNA sequences can generate “AND” logical operations (e.g. EcoRV and SmaI sites in d-DNA); 2) inserting restriction site into i-DNA is equal to a “NOT” operation; 3) multiple restriction sites in one sequence can generate “OR” logical operation. By combining these keys, complex operation can be realized (**Fig. 4e**).

**Figure 4.**
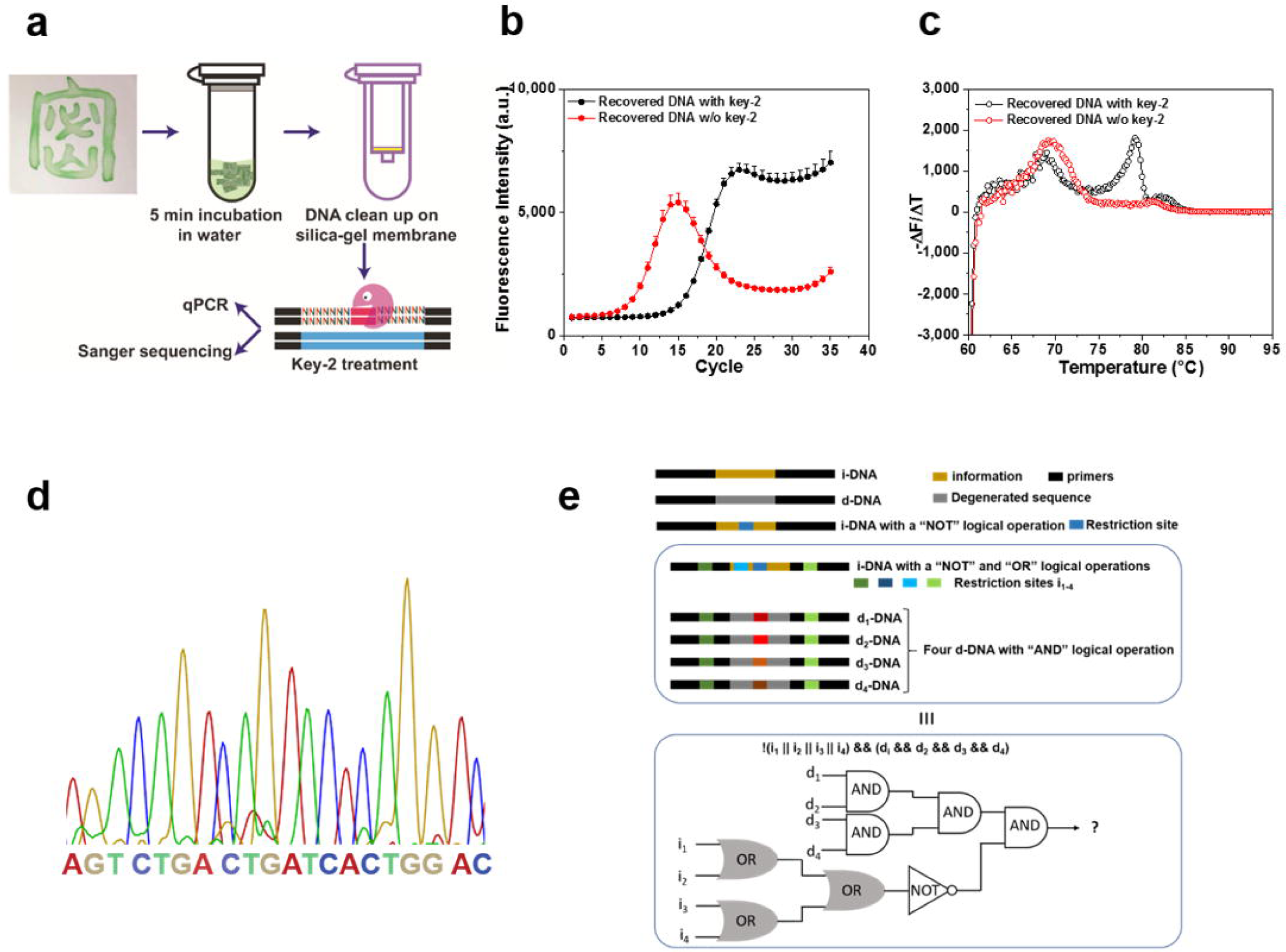
(a) Signature of the ancient Chinese character containing DNA steganography and the workflow of i-DNA recovery. qPCR amplification curves (b) and melting curves (c) of recovered DNA from the signature before and after treatment with key-2. (d) Correct sequence of i-DNA was revealed by PCR and Sanger sequencing. (e) Multiple restriction sites in different d-DNA sequences can generate “AND” logical operations (e.g. EcoRV and SmaI sites in the d-DNA); When restriction site is inserted into any part of i-DNA, it will create a logical operation of “NOT”. When the interceptor uses the wrong key, the information will be destroyed. Multiple restriction sites in one sequence can generate “OR” logical operations, as any one of the restriction enzymes can cleave the sequence. Therefore, by combining these keys, complex logical operation such as “! (i_1_ ‖ i_2_ ‖ … ‖ i_n_) && (d_1_ && d_2_ && … && d_n_)” can be realized. Where “!”, “‖”, “&&” represent NOT, OR, AND logical operations, respectively. i_n_ is a restriction site in i-DNA, while d_n_ is a restriction site in d-DNA.

## Discussion

According to quantum mechanics, sub-atomic particles can simultaneously exist in more than one state and any attempt to detect the particles’ behavior force the wave function to “collapse” in defined state, changing the original particles behavior (Heisenberg Principle). The quantum superposition concept, elegantly illustrated by the Schrödinger’s cat thought experiment, although sometimes considered as a philosophical paradox, has been successfully used for developing QKD-based quantum cryptographic communication networks. In essence, if the message is intercepted, the eavesdropper will leave irreversible traces and can be subsequently detected and alert the receivers that a key has been compromised. A solution of DNA molecules does not possess the property of quantum superposition. However, if an attempt of measurement can change its composition and leave a permanent trace, the design of DNA steganography can be considered as a QKD-like function. When Alice conceals information in a small amount of DNA (e.g., < 1 pmol), PCR is an inevitable step to prepare the sample for sequencing. The incorporation of restriction sites and randomized domains into the DNA masking sequences will result in interesting properties: the heat-generated mismatches cannot be recognized by restriction enzymes, and the difference in diversity after enzyme treatment can be easily detected by a number of different methods (e.g. qPCR melting curve, ΔC_t_, and amplification curve shape). Therefore, without knowing the secondary key (i.e., the nature of the endonuclease) in addition to the correct PCR-amplification primers (the primary key), the message can neither be analyzed nor be copied by eavesdropper (Eve), while Bob (the receiver) can eventually detect the interception.

As the Chinese saying goes: “While the good climb a foot, the wicked climb ten; it takes constant vigilance to stave off evil.” The method to protect information needs always to be discussed in the context of the state-of-the-art technology to read it (supplementary information section **1**: Extended discussion). We have shown a novel DNA steganography utilizing basic molecular biology techniques employing restriction enzyme as a second secret key. Compared to classical DNA steganography only using primers as key, this approach provides a second layer of mask to cover the information, therefore, brings massive complexity against brute-force attack by an eavesdropper. Moreover, this method confers the mechanism to detect interception. Eventually, once the sample is “measured”, it can no longer be processed by enzymes to uncover the information, mimicking a self-destruction feature.

With the advances in different digital and engineering technologies, it has become increasingly easy to forge a physical object, e.g. hand-writing or art piece, indistinguishable from its original. If we can incorporate the DNA steganography as signature into the materials, it will allow the traditional function of signature to meet the challenges from modern technologies. The self-destruction function and QKD-like function can prevent forgers from characterizing and synthesizing the materials, while only the authorized person can confirm the authenticity with the complex keys. While the steganography has demonstrated its utility as encrypted ink for writing, it can also have other applications from personalized art materials to encrypted bank notes.

