## Supplementary material for "Incorporating Randomness into DNA Steganography to Realize Secondary Secret key, Self-destruction, and Quantum Key Distribution-like Function"

**1. Methods**

**Reagents and oligonucleotides**

All oligonucleotides were purchased from IBA life sciences (Göttingen, Germany) in molecular biology grade. Restriction enzymes were purchased from Thermo Fisher Scientific (Waltham, USA). Phusion high-fidelity DNA polymerase was from New England Biolabs and qPCR master mix was from Quantabio Genomics (Beverly, USA). Agarose gel extraction kit was purchased from Qiagen (Venlo, Netherlands). Sanger sequencing was performed by Eurofins Genomics (Ebensburg, Germany).

**Investigation of suitable ratio between i-DNA and d-DNA**

The single stranded i-DNA was mixed with single stranded d-DNA in 1:1, 1:10, and 1: 100 ratios. 5 pmol of i+d-DNA was mixed with excess forward primer and subjected to thermocycling with following protocol. The reaction mixture (50 µL) contained 10x HF buffer, dNTPmix (each 10 nmol), Phusion high-fidelity polymerase (1 U) and template DNA annealed with forward primer. Thermocycling protocol was : 45 s at 98 °C, 5 cycles of 1 min at 55 °C, and 30 s at 72 °C, closing the cycle, final extension for 10 min at 72 °C, and storing at 4°C.

To perform Sanger sequencing, dsDNA product was diluted and subjected to PCR using extended sequencing primers with following protocol: 45 s at 98 °C, then 2 cycles of: 30 s at 98 °C, 1 min at 55 °C, and 30 s at 72 °C, closing the cycle, 18 cycles of: 30 s at 98°C, 1 min at 65 °C, and 30 s at 72 °C, followed by 10 min at 72 °C, and storing at 4°C. Then the reaction mixture was loaded on 2% agarose and 90 V of constant electric field was applied. DNA bands were visualized by a UV transilluminator. The DNA bands of correct size were sliced out and subjected to gel purification using Qiagen gel extraction kit. Purified DNA was mixed with sequencing primer, which annealed to the complementary strand of i-DNA, and Sanger sequencing was performed.

**Preparation of the authentic message by Alice**

1:100 ratio was selected for downstream experiments. Alice sends out 1 pmol of i+d-DNA.

**Eve’s operations upon interception**

The 1:100 i+d-DNA pool was amplified using correct primers (Eve1) and wrong primers (Eve2). The reaction mixture (50 µL) contained 10x HF buffer, dNTPmix (each 10 nmol), Phusion high-fidelity polymerase (1 U) and template DNA (1 pmol), and 50 pmol of forward and reverse primers. PCR was performed with following protocol: 45 s at 98 °C, 25 cycles of 15 s at 98 °C, 1 min at 55 °C, 30 s at 72 °C, close cycle, final extension for 10 min at 72 °C, then store at 4 °C. Sample R contained neither polymerase nor primers and were incubated in the same way as Eves’ DNA.

**Bob’s operations after receiving the DNA**

The DNA samples, that Bob would get, were treated with 10 U of corresponding restriction enzyme in 50 µL reaction volume containing 10 x buffer (cutsmart buffer (NEB) for SmaI or buffer R (Thermo Fisher Scientific) for EcoRV). Enzyme treatments were performed in thermocycler for 1 h (SmaI at 30 °C and EcoRV at 37 °C) followed by 20 min deactivation at 80 °C.

After enzyme treatment, the mixture was diluted 25 times and subjected to qPCR measurement (Thermo Scientific PikoReal Real-Time PCR System) using following protocol. 10 s at 95 °C, 30 -40 cycles of : 15 s at 95°C, 20 s at 55 °C, 30 s at 72 °C, data acquisition, closing the cycle, 30 s at 72°C , then starting melting procedure from 60 °C to 90 °C with holding time 1 s, temperature increment after holding 0.2 °C.

**Decryption of authentic message using complex key-2**

The DNA pool from Alice was incubated with SmaI at 30 °C for 1 h. Next, due to buffer difference and temperature difference, the SmaI digested product was diluted 25 times and PCR amplified for 5 cycles. 5 µL of the PCR product was then incubated with EcoRV at 37 °C for 1h. Each step was monitored by qPCR and Sanger sequencing as described.

**DNA steganography applied as signature**

1 pmol of Alice’s DNA was mixed with water soluble ink. The mixture was then applied on to a cellulose paper (d= 2 cm) and left to dry. To recover the DNA, the paper was sliced and soaked in 300 μL water and incubated for 5 min. Next, the paper pieces were removed and the solution was purified by a DNA purification based on silica membrane. The resulting DNA was then directly subjected to restriction enzyme digestion and subsequent qPCR analysis as well as Sanger sequencing.

**2. Extended discussion**

“While the good climb a foot, the wicked climb ten; it takes constant vigilance to stave off evil.”- Ancient Chinese proverb

Probably there is no truly unbreakable encryption. Herein, we would like to discuss some possible methods, which can be used by an interceptor. As the paper is mainly about the principle of using randomness as a tool in steganography, some relatively trivial details are discussed in this supporting information. We assume that the interceptor aims to read the intercepted message, however, he/she must also pass the message to the intended recipient. Otherwise, the sender and recipient will consider the communication route invalid.

**The message sample can be prepared to prevent the use of deep-sequencing.** The standard preparation of message DNA sample is to mix the synthetic sequence(s) with genomic DNA. The genomic DNA provides a basic mask (as protective layer 0) to prevent the sample from reading using deep-sequencing. If the interceptor has access to key-1 (two primers, as protective layer 1), he/she can amplify the message information specifically. However, this procedure will not work with our steganography design, as there is a second mask (d-DNA, as protective layer 2). Moreover, to amplify the message with correct primers (key-1) without the secondary key will lead to a self-destruction and affect the sample composition (leaving a trace). It is important to note that the randomness associated with genomic DNA is different from the randomness in d-DNA. The genomic DNA will very unlikely form stable duplex with i-DNA, as they share very low sequence similarity, while the d-DNA is designed for such proposal.

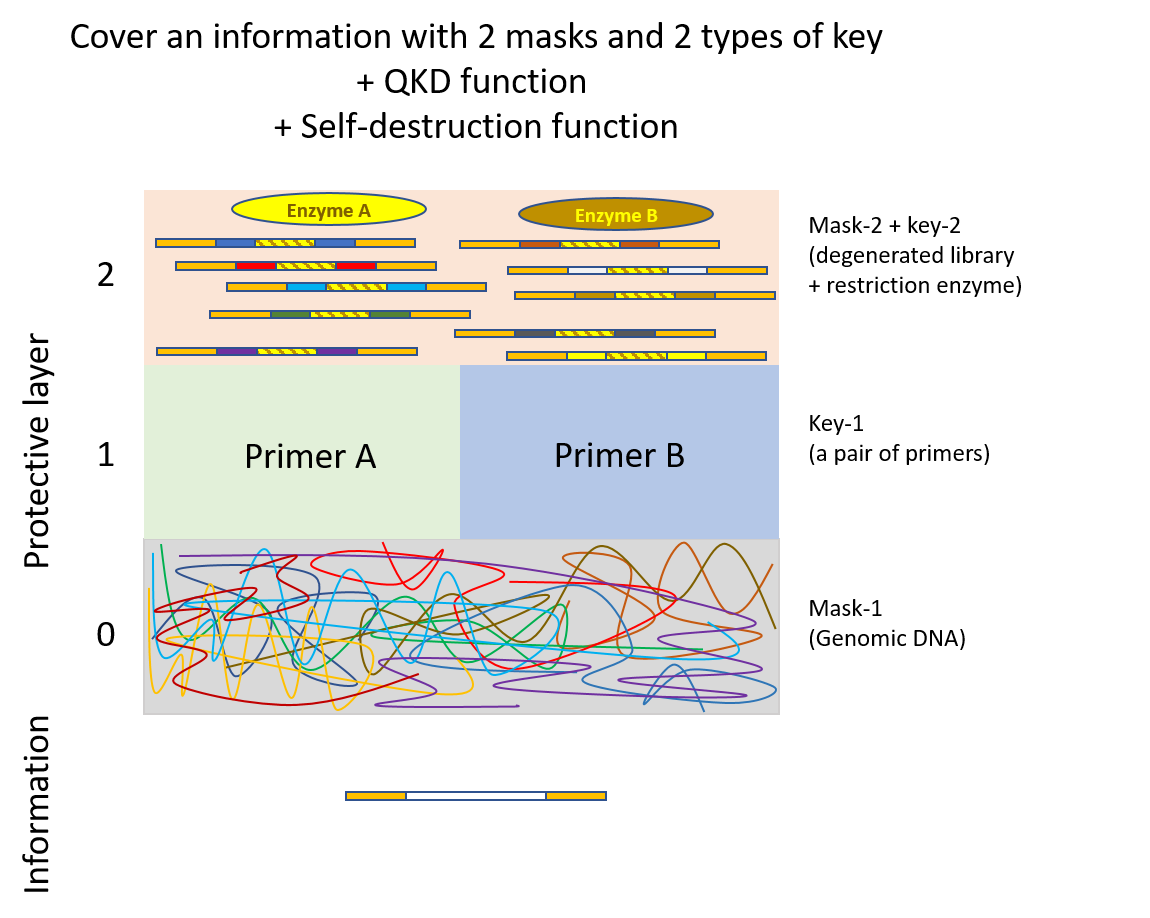

**Scheme S1.** Enhanced DNA steganography with three layers of masks. Layer 0: fragmented genomic DNA as in conventional DNA steganography. Layer 1: Correct primers are required to specifically target i-DNA. Layer 2: degenerated library and corresponding restriction enzyme can protect the message from interceptor with correct primers.

**The message sample is in small quantity, to prevent it from excessive testing.** To amplify an oligonucleotide sample with PCR has its limit. It is well known that extensive PCR multiplexing (e.g. > 40 cycles) will generate large amount of unwanted products, e.g. double stranded DNA that is much larger than the expected size. Moreover, we have discovered that the PCR product will become even more “dirty” when the template is a highly diverse sequence mixture sharing the same primers.^[1]^ Therefore, if the sender prepares the sample in a quantity just sufficient for the recipient to read the message, the interceptor must consume a significant amount of sample, if not the entire sample, for his/her attempts to decrypt the information. To test many different keys (primary keys, secondary keys, and their combinations) will need the message sample in large quantity. Therefore, the amount of sample will make it impossible for the interceptor to make an extensive test, not mentioning to do so with only a neglectable fraction of the original sample.

**3. Supplementary figures**

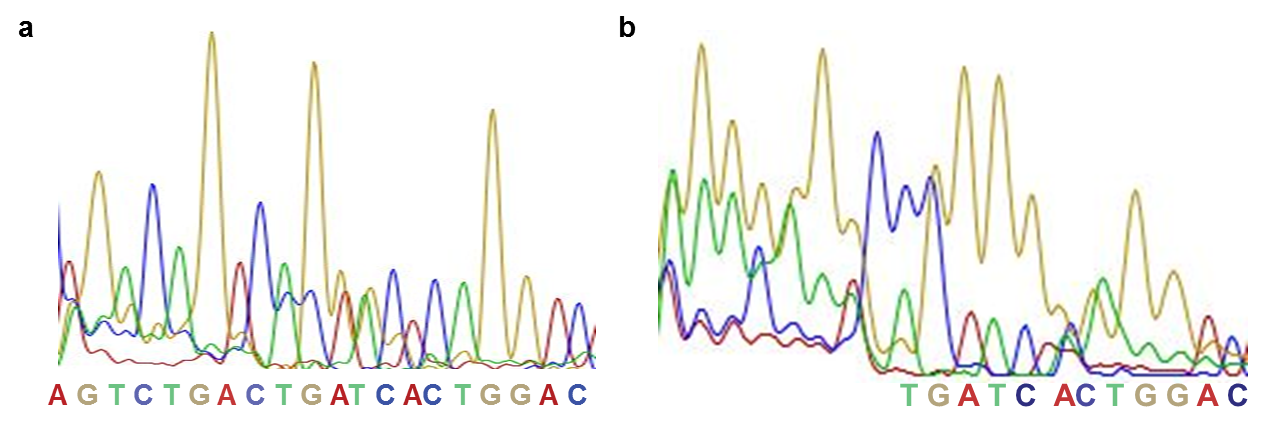

**Figure S1.** Sanger sequencing chromatogram of i+d-DNA at the ratio of 1:1 (a) and 1:10 (b).

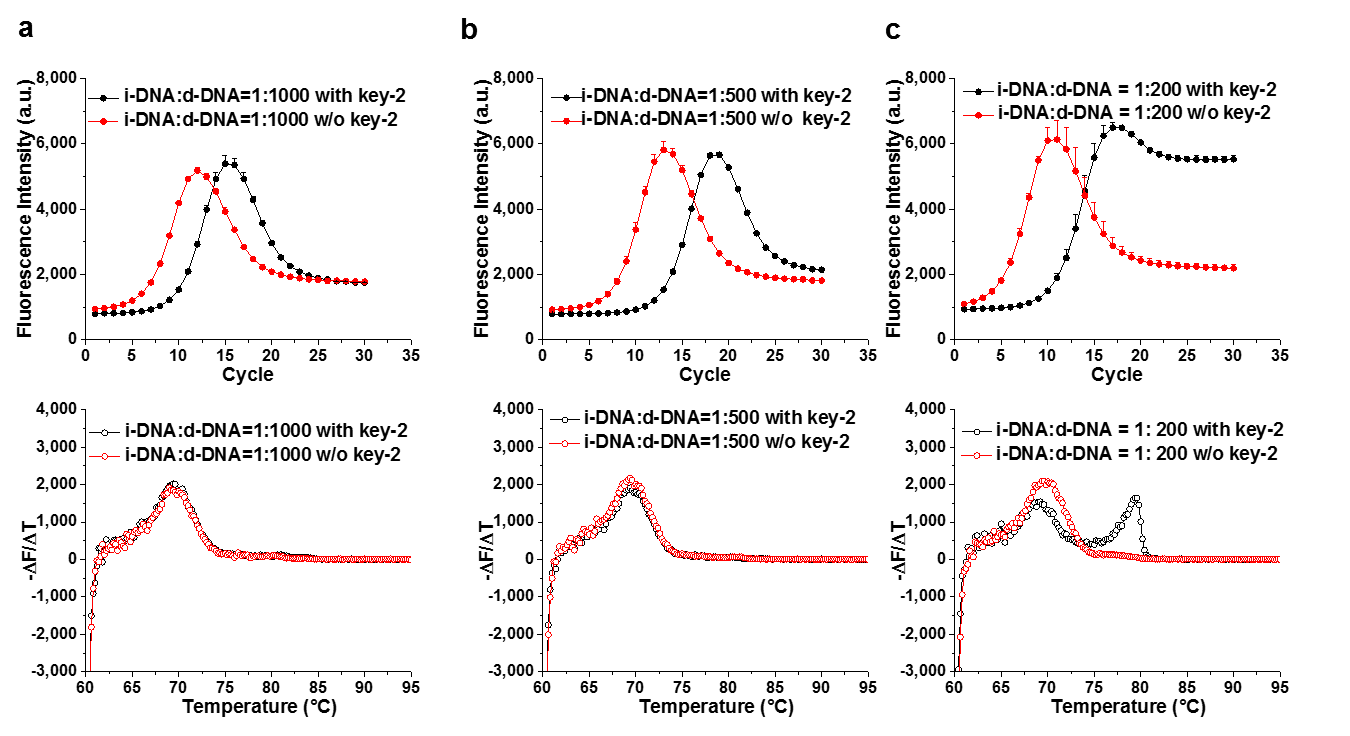

**Figure S2.** qPCR amplification curves (top) and melting curves (bottom) of i+d-DNA at the ratio of 1:1000(a), 1:500(b) , and 1:200(c) with and without key-2 treatment

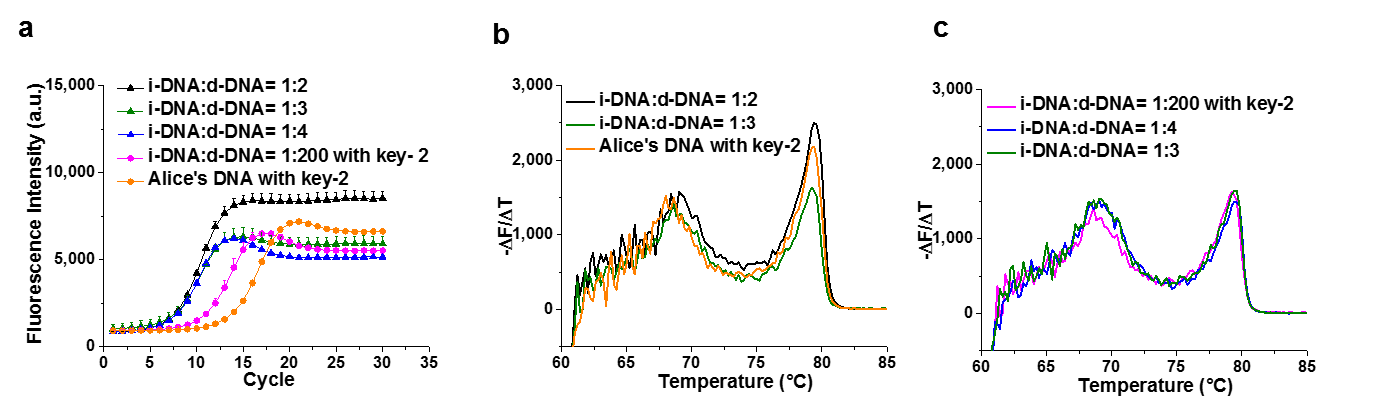

**Figure S3.** Comparison of key-2 treated i+d-DNA (1:200 mixture and 1:100 mixture) with untreated i+d-DNA at the ratio of 1:2, 1:3, and 1:4 via qPCR amplification curves (a) and melting curves (b and c).

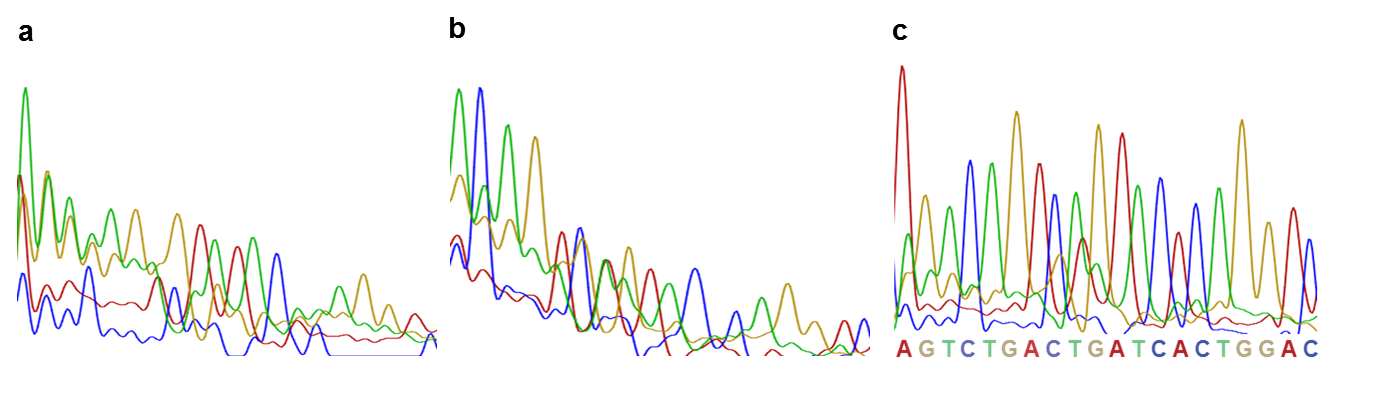

**Figure S4.** Sanger sequencing chromatogram of i+d-DNA at ratio of 1:1000 (a), 1:500 (b), and 1:200 (c) after key-2 treatment.

**4. Supplementary tables**

Table S1 Sequence of oligonucleotides comprising DNA steganography

| Oligonucleotides | Sequence (5’ to 3’) |
| --- | --- |
| i-DNA | GACAATTCACACACGTCCGCAGTCTGACTGATCACTGGACATGAGATCGGAAGAGCGTCG |
| d-DNA with SmaI restriction site | GACAATTCACACACGTCCGC NNNNNNN CCCGGG NNNNNNN ATGAGATCGGAAGAGCGTCG |
| d-DNA with EcoRV restriction site | GACAATTCACACACGTCCGC NNNNNNN GATATC NNNNNNN ATGAGATCGGAAGAGCGTCG |

Table S2 Primer Sequence

| Primers | Sequence (5’ to 3’) |
| --- | --- |
| Desired key-1-R | CGACGCTCTTCCGATCTCAT |
| Desired key-1-F | GACAATTCACACACGTCCGC |
| Wrong key-1-R | GAGATCGGAAGAGCGTCG |
| Wrong key-1-F | TGGTCTCAGCCGCCCTAT |
| PCR primer for Sanger sequencing-R | ACACTCTTTCCCTACCCGACAACCTACTGCCTTTCGAACCTAGACGAC AATTCACACACGTCCGC |
| PCR primer for Sanger sequencing-F | CAAGCAGAAGACGGCATACGAGATCGACGCTCTTCCGATCTCAT |
| Sequencing primer | ACACTCTTTCCCTACCCGACAACCTACTGCCTT |
